# A hybrid machine learning and enzyme-constrained metabolic model for *ab initio* prediction of proteome reallocation

**DOI:** 10.64898/2026.07.20.739489

**Authors:** Ehsan Motamedian, Zoran Nikoloski

## Abstract

High expression of heterologous proteins in microbial cell factories frequently triggers a severe burden due to reallocation of finite cellular proteome. Conventional constraint-based models struggle to predict these resource shifts *ab initio* without relying on condition-specific omics data. To bridge this gap, we developed the Hybrid Transcription-Translation (HyTT) framework, combining multivariate adaptive regression splines (MARS) with enzyme-constrained metabolic models by enforcing an 80S ribosome integrity constraint. Cast as a mixed-integer linear programming problem, HyTT mathematically couples macroscopic spatial boundaries with microscopic, sequence-derived translational costs based on a bisection search. Validation against steady-state chemostat quantitative proteomics data demonstrated the superior capability of HyTT over contenders in predicting system-wide resource (re)allocation in *Saccharomyces cerevisiae*. Operating *ab initio*, the framework doubled the predictive accuracy of protein abundances (Pearson r=0.501) compared to conventional models, successfully segregating the minimal essential proteome from the cellular reserve pool. Crucially, HyTT autonomously captures complex stress responses vital for metabolic engineering. Upon simulating a 15% recombinant protein burden, the framework accurately predicted systemic growth retardation, decrease of ribosomal portion of the proteome, and surge of ethanol production, in line with the Crabtree effect. System-level analysis uncovered that cells adapt to restricted proteomic capacity through non-uniform metabolic rerouting, downregulating respiratory complexes in favor of high-turnover glycolytic enzymes, and relying on ribosomal paralog switching to minimize sequence-specific assembly costs. Ultimately, HyTT provides a computationally agile, sequence-driven platform for decoding dynamic resource reallocation, offering a powerful predictive tool to navigate metabolic trade-offs and guide rational strain design without requiring condition-specific multi-omics inputs.

## 1. Introduction

*Saccharomyces cerevisiae* serves as the principal eukaryotic cell factory for industrial biosynthesis of high-value chemicals, biofuels, and therapeutic proteins. However, the high expression of synthetic constructs and heterologous pathways invariably imposes a severe physiological penalty on the host, widely recognized as metabolic burden (Fujita et al., 2024). Recent advances in systems biology and multi-omics profiling have revealed that this metabolic burden extends far beyond energetic depletion, such as competitive ATP and GTP hydrolysis, and is rather driven by resource allocation constraints, widely recognized as protein burden (Kafri et al., 2016). During acute heterologous expression, recombinant proteins physically sequester a substantial fraction of the finite cellular proteome, impairing the functionality of the endogenous metabolic network and the inherently massive translation machinery. Contemporary cytological studies have demonstrated that the resulting protein burden triggers profound systemic stress responses, including: defective nucleolus formation, an imbalance in ribosomal RNA (rRNA) biogenesis, and induction of nitrogen starvation-like states regulated via the inactivation of the TORC1 pathway (Tartik, 2026). To alleviate this resource allocation constraint, fluxes are redistributed so highly efficient but space-consuming respiratory pathways are downregulated in favor of compact, high-flux fermentative routes, ultimately driving the onset of overflow metabolism and macroscopic growth retardation (Eguchi et al., 2018).

To quantitatively predict these systemic resource reallocations and guide stress-aware rational strain design, constraint-based reconstruction and analysis (COBRA) methodologies have devised and applied a set of advanced computational tools (Orth et al., 2010). Whilst traditional genome-scale metabolic models (GEMs) compute flux balances based on stoichiometry and thermodynamics, they inherently fail to capture physiological phase shifts induced by spatial macromolecular crowding. To bridge this gap, enzyme-constrained models (ecGEMs), such as the pioneering GECKO framework (Domenzain et al., 2022), revolutionized phenotype predictions by integrating absolute quantitative proteomics, molecular weights, and enzymatic turnover numbers (kcat) to strictly bound the allowable physical proteome space. The evolution of these models has been remarkably rapid; the recent introduction of GECKO 3.0 successfully integrated deep learning models (e.g., DLKcat) to autonomously predict global catalytic efficiencies (kcat) directly from amino acid sequences, thereby drastically expanding predictive capabilities in the absence of experimental kinetic data (Chen et al., 2024).

Despite these computational advancements, ecGEMs estimate the fundamental metabolic synthesis cost of a target protein based exclusively on its linear polypeptide length and raw molecular mass. This simplification systematically neglects the effect of mRNA sequence features such as codon adaptation indices, minimum free energy of untranslated regions, and specific amino acid stoichiometries on active translational kinetics (Verhagen et al., 2026). Recent biophysical evidence in *S. cerevisiae* has unequivocally demonstrated that evolutionary selection pressure does not merely optimize generic translation efficiency, but rather strictly optimizes ribosome utilization (Sharma, 2023). Because the biogenesis of the ribosomal complex consumes massively more cellular resources than transcription, the substantial metabolic penalty of a recombinant sequence is dictated by its ribosome utilization efficiency. Consequently, models lacking sequence-specific translational constraints fail to capture inherent translational costs, leading to artificial parity between fundamentally distinct metabolic pathways.

In parallel efforts, Resource Balance Analysis (RBA) models and comprehensive Metabolism and macromolecular Expression models (ME-models) have been developed to explicitly account for both the transcription and translation machineries. While yeast-specific RBA models (e.g., scRBA) (Dinh and Maranas, 2023) and ME-models offer a highly detailed stoichiometric accounting of cellular self-replication (O’Brien et al., 2013), they remain computationally challenging due to extensive condition-specific kinetic parameters and the inherent non-linearity of explicitly modeling massive sequence data spaces (Lloyd, 2018). Consequently, these models remain rigid when attempting to simulate the dynamic ribosome sequestration and targeted metabolic redistribution characteristic of an acute recombinant protein burden.

Bridging this mechanistic gap requires a paradigm shift: a framework capable of translating complex, sequence-based translational kinetics into solvable, dynamic physical constraints, without incurring the nonlinear computational overhead inherent to full ME-models. Machine Learning (ML) has emerged as a transformative bridge in this context, demonstrating an unparalleled capability to extract hidden biophysical regulatory grammar from expansive multi-omics datasets and map physicochemical sequence architectures directly to functional translation efficiencies (Cuperus et al., 2017). By embedding these sequence-derived predictions within stoichiometric metabolic networks, it becomes possible to mathematically formalize sequence-dependent translational costs and force the network to navigate a strictly constrained proteomic and bioenergetic landscape.

In this study, we present the Hybrid Transcription-Translation (HyTT) framework, a fully *ab initio* computational platform that integrates robust enzyme-constrained metabolic modeling with sequence-driven machine learning predictions. By embedding sequence-specific Multivariate Adaptive Regression Splines (MARS) penalties as constraints in the optimization problem, HyTT autonomously couples intricate translational costs to global stoichiometric limits via mixed-integer linear programming (MILP). Through comprehensive simulations, we demonstrate the unique capability of HyTT to mathematically decouple the highly flexible metabolic proteome from the inherently rigid translational machinery, and, critically, to distinguish between the active catalytic proteome and the inactive reserve pool (the idle proteome). Upon simulating a 15% heterologous burden, HyTT successfully predicts the emergence of the Crabtree effect *ab initio*, recapitulating the reduction of the ribosomal proteome fraction recently observed *in vivo* (Fujita et al., 2024; Metzl-Raz et al., 2017). Furthermore, the framework uncovers sequence-driven metabolic redistribution across core pathways, highlighted by the targeted paralog and isoenzyme switching required to minimize sequence-specific translational costs within a restricted proteomic capacity. By actively evaluating the biophysical costs of recombinant expression, the HyTT framework provides a versatile computational platform that, once trained on a single reference state, predicts resource reallocation in unseen conditions without requiring additional condition-specific omics data. Finally, by precisely quantifying the resulting protein burden, HyTT offers a robust foundation for next-generation rational strain design and metabolic engineering.

## 2. Materials and Methods

### 2.1. Data collection and definition of target variable

To train the machine learning component, enabling the extraction of fundamental sequence-dependent translational rules, the framework initially utilizes a high-quality baseline multi-omics dataset. It is crucial to note that this dataset is employed exclusively for the initial model training; once the sequence-to-expression mapping is learned, the HyTT framework operates entirely *ab initi*o and does not require condition-specific omics constraints for subsequent metabolic simulations. Absolute quantitative data for protein and mRNA abundances under multiple physiological conditions were obtained from the comprehensive study of *S. cerevisiae* by Lahtvee et al. (Lahtvee et al., 2017). The final dataset compiled for this study consisted of 1,786 well-characterized genes. To establish a robust composite measure of cellular resource commitment and protein expression efficiency, the target variable was defined as P × L (Protein abundance × Protein length), calculated in units of mmol/gDW × number of amino acids. Unlike simple abundance metrics, the P × L index accurately reflects the total translational resource allocation (i.e., the absolute translational burden) required by the cell to maintain the functional pool of each specific protein.

### 2.2. Feature engineering and considered biological layers

To capture the thermodynamic and biophysical rules governing protein expression, an extensive feature engineering pipeline yielded 115 numerical predictors. Features were extracted using custom Python (Biopython (Cock et al., 2009)) and R (seqinr) scripts and functionally categorized into four biological layers:

#### Transcriptional and mRNA Stability

Captures transcript availability and maintenance costs using baseline mRNA abundance (M), coding sequence (CDS) length, experimentally corrected mRNA half-lives, and mRNA oxidative potential (CDS guanine frequency).

#### Translational Regulation

Dictates decoding efficiency via the Codon Adaptation Index (CAI), Kozak scores, and UTR thermodynamic barriers (Minimum and Ensemble Free Energies). Specific CDS nucleotide percentages (e.g., AT, A, and AA) were included to evaluate secondary structure and elongation speed.

#### Post-Translational and Protein Stability

Governs functional lifespan using protein half-lives, instability indices, isoelectric point (pI), GRAVY scores, and structural disordered fractions. To quantify ROS vulnerability, we engineered the Protein Oxidative Potential (length-normalized cysteine frequency) and its ratio to the mRNA oxidative potential.

#### Evolutionary and Compositional

Evaluates basal sequence architecture through total transcript/protein lengths, evolutionary signatures, and specific amino acid frequencies, discriminating between metabolically expensive and inexpensive residues.

### 2.3. Data preprocessing and partitioning

Prior to model training, the dataset underwent rigorous preprocessing. Non-numerical attributes were excluded, and genes with missing values (NAs) were omitted, resulting in a robust, complete matrix of 115 features for 1710 proteins (see Supplementary file 1 for a comprehensive glossary of all extracted features, categorized by their respective biological layers alongside their calculated values). To ensure the machine learning model could be properly generalized and evaluated against unseen data, the dataset was split into a training set (80%) and a hold-out test set (20%). This partitioning was performed using a stratified sampling approach based on the target variable (P × L), utilizing the createDataPartition function from the caret package in R. This stratification guaranteed that the underlying distribution of translational efficiencies was uniformly preserved across both training and testing sets, effectively mitigating data imbalance and preventing model overfitting.

### 2.4. Multivariate Adaptive Regression Splines (MARS) implementation

To construct a highly interpretable, non-parametric model to predict absolute protein abundances and sequence-specific translational costs, Multivariate Adaptive Regression Splines (MARS) was employed using the earth package in R. The algorithm evaluated 115 comprehensive genomic and transcriptomic features to predict the sequence-specific Translational Burden (P × L), defined as the product of protein expression and protein length, ensuring that the derived resource allocation rules were robustly applicable across the *S. cerevisiae* proteome. Unlike black-box algorithms (e.g., Random Forests or Deep Neural Networks), MARS generates an explicit mathematical formulation consisting of hinge functions (piecewise linear segments), making it uniquely suited for seamless integration into the linear constraints of GEMs.

To ensure strict biological interpretability and prevent mathematical overfitting, the model hyperparameters were meticulously constrained. The maximum degree of interaction was restricted to 1 (degree = 1), enforcing an additive model architecture where the independent contribution of each biological layer could be individually assessed. To heavily penalize model complexity and promote feature parsimony, the generalized cross-validation (GCV) penalty was set to a high value (penalty = 10), with a maximum of 30 retained terms (nprune = 30). Model robustness and generalization were rigorously validated using a 10-fold cross-validation scheme repeated 30 times (nfold = 10, ncross = 30).

### 2.5. Formulation of the HyTT framework

To bridge the gap between purely stoichiometric Flux Balance Analysis (FBA) and sequence-dependent proteome allocation, we developed the Hybrid Transcription-Translation Constrained Metabolic Model (HyTT). This framework effectively transforms a static metabolic network into a system limited by the physical and sequence-based constraints.

#### 2.5.1. Baseline enzyme-constrained model and network expansion

The enzyme-constrained genome-scale metabolic model of *S*. *cerevisiae* (ecYeastGEM), derived from the consensus yeastGEM v8.6.2 network, served as the stoichiometric foundation for this study. The pre-constructed baseline ecModel was directly loaded via the GECKO 3 toolbox environment (Chen et al., 2024), utilizing its pre-integrated, high-quality, and manually curated k_cat_ values. The native reconstruction contains 1,160 metabolic genes with pre-assigned enzymatic functions.

To enable the HyTT framework’s multi-layered constraints, we first deactivated the standard GECKO enzyme-usage formulations by constraining the upper bounds of all default usage_prot_ pseudo-reactions to zero. Subsequently, the baseline metabolic network was expanded by introducing a unified set of protein-synthesis pseudo-reactions (ml_prot_, denoting machine learning-coupled protein synthesis) for both endogenous metabolic enzymes and the 137 core ribosomal subunits. These newly implemented ml_prot_ reactions serve as the explicit mathematical bridge between the metabolic network and the machine learning layer. Specifically, each reaction dynamically couples the flux of a metabolic or ribosomal protein to its sequence-specific translational cost predicted by the MARS model, thereby enforcing absolute spatial and biophysical boundaries directly on the total proteome allocation. Through these unified ml_prot_ reactions, the expanded network (comprising 1,297 genes in total) dynamically tracks the exact synthesized proteome mass. This mathematical formulation ensures that the combined physical burden of all active metabolic enzymes and the newly integrated translational machinery strictly adheres to the total cellular protein capacity constraint (P_total_).

Second, to comprehensively account for the transcriptomic footprint, a dedicated intracellular mRNA pool (mRNA_pool[c]) and corresponding exchange reactions were introduced. To ensure a consistent nomenclature and facilitate a direct one-to-one stoichiometric mapping between the transcriptome and the proteome, each mRNA species and its associated usage reaction were identified using the UniProt accession number of the corresponding protein (e.g., ml_mRNA_uniprot). This modification allows the model to process true biological molecular counts, enabling an integrated representation of transcriptional and translational constraints within the HyTT framework. To ensure steady-state feasibility under strict coupling constraints, standard sink reactions for transcripts and proteins were implemented to simulate continuous turnover and prevent algorithmic dead-ends (details in Supplementary file 2, Note S1).

Crucially, while the expanded stoichiometric network comprises 1,297 genes (1,160 metabolic and 137 ribosomal), the MARS algorithm was trained on a broader dataset of 1,786 genes (Section 2.4). This inclusive training landscape allows HyTT to capture universal, system-wide translational rules, ensuring the sequence-derived constraints are biologically robust and avoid overfitting to specific metabolic pathways. The calculated biophysical features and ML-derived parameters for the 1,297 mapped network genes are provided in Supplementary file 3.

#### 2.5.2. Mathematical formulation and biophysical constraints

To accurately simulate the metabolic burden of recombinant protein production and dynamic cellular resource reallocation in *S. cerevisiae*, the HyTT framework was formulated as a penalty-driven MILP problem. This hybrid mechanistic-statistical model integrates the stoichiometric constraints of the GEM with dynamic enzyme capacities and sequence-dependent translational penalties. Unlike traditional data-forced models that impose rigid upper bounds on enzyme fluxes, HyTT utilizes sequence-derived translational and transcriptional costs as dynamic penalties, allowing the network to autonomously identify the most resource-efficient metabolic routing. The optimization problem and its associated biophysical parameters are defined as follows:

##### Objective Function

The primary biological objective is the maximization of the specific growth rate (μ):

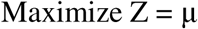

##### Mass balance and thermodynamic constraints

Intracellular metabolites are assumed to be in a pseudo-steady state, where metabolic fluxes (v) are constrained by the stoichiometric matrix (S):

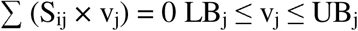

##### Enzymatic capacity and exact integration of MARS

For the 1,160 integrated metabolic genes, the maximum reaction flux is bounded by the enzyme concentration ([E]_j_), the catalytic turnover number (k_cat,j_), and the in vivo saturation factor (σ):

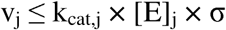

To accurately reflect the sub-optimal thermodynamic conditions within the cell, the global saturation factor (σ) was calibrated to 0.7 (Domenzain et al., 2022). The MARS algorithm predicts the total translational resource allocation (pl_j_ = P × L) using a linear combination of hinge functions (H_k_), integrated via the Big-M method:

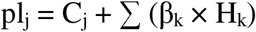

where H_k_ = max(0, x_k_ - c_k_). The linearization is enforced using binary variables (z_k_ ∈ {0, 1}) and a sufficiently large constant (M):

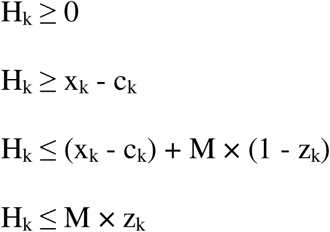

##### Global Proteome Allocation

The total mass of metabolic enzymes, ribosomes ([P]_ribo_), and, when applicable, the recombinant product ([P]_recomb_) is constrained by the effectively modeled fraction (f) of the total protein capacity (P_total_):

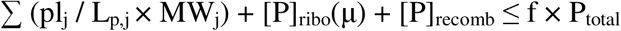

Based on comprehensive quantitative proteomics, the total cellular protein capacity (P_total_) was defined as 0.46 g/gDW (Lahtvee et al., 2017). Furthermore, the modeled proteome fraction (f) was set to 0.71, explicitly accounting for the ∼9% mass contribution of the incorporated core ribosomal machinery estimated from (Lahtvee et al., 2017).

##### Translational capacity and growth laws

The protein synthesis rate is limited by the translation elongation rate (k_trans_) and the active ribosome concentration ([R]_active_), accounting for both growth (μ) and continuous protein degradation (k_deg,p_):

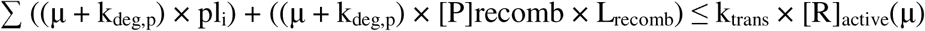

To ensure strict biophysical realism, the translation elongation rate (k_trans_) was set to 28,800 amino acids/h (von der Haar, 2008), and the average protein degradation rate (k_deg,p_) was defined as 0.05 1/h (Christiano et al., 2014). The active ribosome pool is dynamically estimated using phenomenological growth laws:

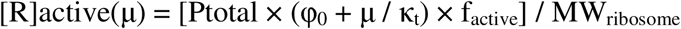

The parameters for this dynamic estimation were calibrated as follows: the baseline ribosomal fraction (φ_0_) was set to 0.05, and the translational efficiency constant (κ_t_) was defined as 1.3 1/h (Metzl-Raz et al., 2017). Additionally, the active ribosomal fraction (f_active_) was set to 0.85 (von der Haar, 2008), and the molecular weight of the eukaryotic 80S ribosome (MW_ribosome_) was established at 3 × 10^6^ Da (Ben-Shem et al., 2011).

##### Transcriptional capacity

The total rate of mRNA synthesis must cover dilution by cell division and continuous transcript degradation (k_deg,m_), bounded by the RNA polymerase capacity ([RNAP]_total_) and the transcription elongation rate (k_poly_):

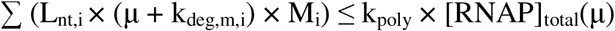

For the transcriptional constraints, the elongation rate (k_poly_) was set to 180,000 nucleotides/h (Mason and Struhl, 2005), and the average mRNA degradation rate (k_deg,m_) was defined as 2.1 1/h (Presnyak et al., 2015). The total RNA polymerase capacity ([RNAP]_total_) was estimated at 0.01 g/gDW.

Furthermore, to address the additive machine learning loophole, where highly optimized genes with superior CAI might theoretically generate positive protein flux solely from the MARS intercept without requiring an mRNA template, a biophysical template lock was implemented. To mathematically enforce direct mRNA-protein coupling and explicitly close this algorithmic loophole, the translational concentration (P_prot,j_) was strictly bounded by the magnitude of its corresponding transcriptomic usage abundance (M_mRNA,j_) using a sufficiently large positive constant (M):

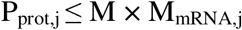

This "ultimate lock" operates as a definitive biophysical constraint: if the solver allocates zero transcriptomic budget to a specific gene (M_mRNA,j_ = 0), the maximum allowable protein synthesis rate is simultaneously and forcibly reduced to zero. This mathematical formulation completely forbids template-free translation, strictly binding the MARS predictions to the physical presence of the transcript.

##### Strict ribosomal stoichiometry

Unlike standard resource allocation models that treat the ribosomal pool as an unconstrained aggregate of independent monomers, the HyTT framework enforces a strict biophysical stoichiometry for the 80S ribosome. In *S. cerevisiae*, the 80S ribosome is a massive 3 MDa complex. To prevent the MILP solver from artificially discarding high-molecular-weight or kinetically expensive ribosomal subunits to minimize the objective function, an algorithmic artifact common in standard parsimony approaches, a strict assembly constraint was integrated into the stoichiometric matrix. A pseudometabolite representing the fully assembled ribosome (P_80S_) was defined. The model mathematically enforces a 1:1 molar ratio for all core ribosomal protein families, and a 2:1 molar ratio for the acidic P-stalk components (e.g., RPP1 and RPP2), which function as dimers in the heteropentameric structure. Furthermore, for ribosomal families possessing whole-genome duplication (WGD) paralogs (e.g., RPL1A and RPL1B), the sum of the molar fluxes of the paralogs was constrained to meet the exact stoichiometric demand of the P_80S_ assembly node. This rigid coupling ensures that the simulated cell pays the exact thermodynamic, spatial, and translational cost of complete ribosome biosynthesis.

#### 2.5.3. Numerical optimization strategy and penalty-driven allocation

The solution of the HyTT framework follows a two-step optimization strategy to resolve the non-linear coupling between the growth rate (μ) and the resource constraints. Because the growth rate functions as a direct multiplier for macromolecular dilution and synthesis boundaries, the resulting formulation is fundamentally a non-linear programming (NLP) problem. To render this computationally tractable at the genome-scale, we employed a strategy widely validated in robust ME models and RBA frameworks.

First, a bisection-based binary search, analogous to the algorithms used in nonlinear ME models (Yang et al., 2016) and iterative scRBA approaches (Dinh and Maranas, 2023), is employed to identify the maximum feasible growth rate within a numerical tolerance of 0.001 1/h. Second, because constraint-based models inherently suffer from alternative optima and degeneracy at a fixed maximum growth rate, the model performs a penalty-driven parsimonious allocation. Similar to the principles underlying parsimonious FBA (pFBA) (Lewis et al., 2010), the MILP objective function is reformulated to minimize the global metabolic and translational cost. Specifically, a negative coefficient is assigned to the global protein pool usage reaction, effectively forcing the solver to minimize the total continuous protein mass required to sustain the identified growth rate (L1-norm parsimony). This is supplemented by a hierarchical penalty system designed to maintain both biological fidelity and numerical stability. A moderate penalty (w = 100) is applied strictly to the protein sink reactions, enabling them to act as algorithmic escape valves without promoting wasteful accumulation. This sequential optimization explicitly breaks mathematical symmetry, forcing the solver to adopt the most biologically realistic and resource-efficient metabolic routing within the restricted spatial capacity.

#### 2.5.4. Integration of recombinant protein synthesis and metabolic burden

To evaluate the HyTT framework under metabolic engineering scenarios, we incorporated heterologous production of Green Fluorescent Protein (GFP). Unlike standard approaches that treat recombinant production as a static external sink, HyTT models the induced stress through a three-layered burden mechanism: precursor demand, spatial burden, and translational burden.

First, precursor demand is captured via an explicit synthesis reaction (r_GFP_synthesis), rigorously accounting for amino acid stoichiometry and the ATP/GTP costs of peptide synthesis (Supplementary file 2, Note S2). Second, the spatial burden is simulated by sequestering a target fraction of the total proteome capacity (P_total_). This mechanism reduces the available baseline space (f_pool_), forcing the endogenous network and ribosomes to compete for survival within a diminished physical volume. Third, the translational burden accounts for kinetic drag caused by heterologous transcripts. We compute an effective global translation rate (κ_t,effective_) as a weighted harmonic mean of host and recombinant translation rates, simulating how suboptimal codons sequester ribosomes and retard global machinery:

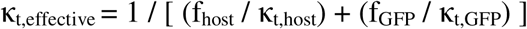

To test HyTT, we simulated an acute stress scenario by pushing GFP production to 15% of total proteome, an experimentally validated tipping point for severe translational constraints in *S. cerevisiae* (Eguchi et al., 2018). Combined with high-flux glucose intake (11.1 mmol/gDW/h), this dual-pressure design forces the cell to accommodate maximal carbon flux and extreme expression, allowing HyTT to predict emergent physiological reallocations, such as the shift from respiratory to hyper-fermentative pathways.

### 2.6. Computational implementation and software environment

While feature engineering and machine learning model development were implemented in Python and R (Section 2.2), all subsequent genome-scale network manipulations, data processing, and metabolic simulations were executed in MATLAB. Specifically, the COBRA (Heirendt et al., 2019) and RAVEN (Wang et al., 2018) Toolboxes were utilized for FBA, and the GECKO 3.0 toolbox (Chen et al., 2024) facilitated sequence-specific kinetic parameter extraction and base enzyme-constrained model reconstruction. The resulting MILP problems were strictly solved using the Gurobi Optimizer.

To benchmark the HyTT framework, two baseline models were implemented: a Basic GECKO (governed solely by a global pool limit and solved via L1-norm parsimony) and a Protein-constrained GECKO (pcGECKO) integrated with absolute proteomics from (Lahtvee et al., 2017). For both baselines, the modeled proteome fraction was set to f = 0.62, representing the cumulative mass of mapped metabolic enzymes from the experimental dataset (Lahtvee et al., 2017). In contrast, HyTT utilizes an expanded fraction of f = 0.71 to incorporate the additional ∼9% mass contribution of the core ribosomal machinery. All models used a global saturation factor of σ = 0.7 to ensure an unbiased comparison of HyTT’s autonomous reallocation capabilities against standard methods. The custom MATLAB scripts, including the MILP architecture construction (buildHyTT.m) and solving algorithms (solveHyTT.m), are openly available at https://github.com/ehsanmotamedian/HyTT.

When evaluating spatial predictive accuracy, a Paralog Aggregation strategy was employed at the proteomic level. Due to the deterministic winner-takes-all nature of linear programming, the MILP solver dynamically selects the most translationally efficient paralog (e.g., highest CAI, lowest MARS cost) and entirely represses suboptimal paralogs. Biological systems, however, exhibit intrinsic regulatory noise leading to concurrent paralog expression. Because whole-genome duplication (WGD) paralogs (e.g., RPL1A and RPL1B) are functionally equivalent and occupy the exact same structural subunit within the 80S ribosome, evaluating them in isolation artificially deflates correlation metrics. Therefore, simulated and experimental abundances of paralogs from the same structural family were aggregated into a single functional abundance prior to calculating Pearson and Spearman correlations. This ensures a biophysically meaningful comparison based on total structural capacity required to assemble multimeric complexes, rather than penalizing the model for theoretical isoenzyme optimization.

## 3. Results and Discussion

### 3.1. *Ab initio* derivation of the translational allocation equation

The primary objective of the HyTT framework is to establish a sequence-driven mapping of translational efficiency that eliminates reliance on *a priori* condition-specific omics data, allowing the integrated metabolic network to dynamically adapt to simulated environmental perturbations. By training a strictly penalized MARS model on a comprehensive multi-omics dataset, we distilled 115 biophysical features into a parsimonious set of 19 basis functions driven by 12 distinct predictors. The model achieved an R² of 0.674 on the training set and 0.672 on a 20% hold-out test set, with a low root mean square error (RMSE ∼ 0.01), confirming robust generalization without overfitting.

The twelve features autonomously selected by MARS reflect core mechanistic checkpoints of protein synthesis. For the transcriptional layer, baseline transcript abundance (M) and the total transcript pool (M × L_m_, where L_m_ is coding sequence length) emerged as the dominant predictors of the steady-state protein abundance × protein length, P×L. Incorporating transcript length directly accounts for the kinetic burden of translation, as longer transcripts prolong ribosome occupation and reduce re-initiation efficiency. Crucially, MARS piecewise linear hinge functions naturally capture biological saturation; unlike rigid linear models, MARS identified that at elevated mRNA levels, translation rates plateau as free ribosomes become limiting, mirroring real physical constraints (Li et al., 2014; Samih et al., 2025).

Decoding efficiency and initiation bottlenecks were mapped via the Codon Adaptation Index (CAI) and specific sequence compositions (CDS-AT, CDS-A, and CDS-AA percentages), aligning with paradigms where elongation speed and transcript secondary structure dictate ribosome stalling (Plotkin and Kudla, 2011). Evolutionary metabolic costs were captured through amino acid frequencies (i.e., alanine, methionine, and leucine), penalizing energetically expensive residues that require heavy ATP and reducing equivalent investments (Akashi and Gojobori, 2002). Finally, post-translational stability and oxidative vulnerability were identified as critical regulatory layers. Through engineered metrics, Protein Oxidative Potential (Cys_len, length-normalized cysteine frequency) and the Oxidative Ratio (Ratio_Ox), the algorithm applies numerical penalties to proteins prone to reactive oxygen species (ROS) degradation, capturing the compounded risk of unstable transcripts translating fragile products (Cabiscol et al., 2000). Collectively, this pipeline generated an explicit, sequence-driven regression equation ready to be embedded as a biophysical constraint within the stoichiometric matrix.

#### Mathematical formulation of the translational burden

Unlike opaque machine learning architectures, MARS yields an explicit, additive mathematical equation easily integrated into linear stoichiometric matrices. The derived universal equation for *S. cerevisiae* computes the predicted translational burden, P × L, as a baseline intercept modified by piecewise linear hinge functions (max(0, x - c) or max(0, c - x)): P × L = 0.0594 + (1.44E-05 × max(0, [M × L_m_] - 8104.54)) - (1.48E-05 × max(0, [M × L_m_] - 9011.82)) - (0.00177 × max(0, 6.467 - M)) + (0.00158 × max(0, M - 6.467)) - (0.0572 × max(0, CDS_AT - 4.475)) + (0.0423 × max(0, CDS_AT - 4.669)) - (0.0144 × max(0, 7.349 - CDS_AT)) + (0.0151 × max(0, CDS_AT - 7.349)) + (0.00109 × max(0, 29.63 - CDS_A)) - (0.00158 × max(0, 10.09 - CDS_AA)) + (0.0287 × max(0, CAI - 0.816)) + (0.2429 × max(0, 0.020 - Ratio_Ox)) + (0.00103 × max(0, A_pct - 7.746)) - (0.00235 × max(0, A_pct - 10.595)) - (0.00052 × max(0, 8.333 - L_pct)) + (0.0148 × max(0, 0.645 - M_pct)) - (3.1751 × max(0, 0.0018 - Cys_len)) + (0.00037 × max(0, 7.20 - HL_Ratio))

The equation uncovers critical biological thresholds. The total transcript pool (M × Lm) triggers structural inflections at ∼8104 and ∼9011, bounding incremental changes in translational burden. Baseline transcript abundance (M) exhibits a sharp inflection at 6.467 molecules/pgDW. Translational efficiency is heavily amplified by optimal codon bias, where CAI values > 0.816 boost capacity via a significant positive coefficient (+0.0287), avoiding ribosome queuing.

Conversely, the model penalizes structural and oxidative vulnerabilities; a high cysteine content (Cys_len) triggers an aggressive penalty (−3.1751) below specific stability thresholds, mathematically capturing the bioenergetic costs of deploying chaperones to refold or degrade fragile proteins under stress (Cabiscol et al., 2000; Tartik, 2026). Lastly, amino acid budgets dictate synthesis bounds, evidenced by tightly constrained thresholds for Alanine (A_pct between 7.7% and 10.6%) and Leucine (L_pct). This transparent rule set demonstrates how static genomic sequences dictate the physical allocation limits of the translational machinery.

### 3.2. Benchmarking predictive performance: Autonomous reallocation versus data relaxation

To evaluate the performance of HyTT, we benchmarked it against Basic GECKO (governed solely by a global pool limit) and proteomics-constrained GECKO (pcGECKO), both sharing a base protein capacity constraint. Under high-glucose conditions (11.1 mmol/gDW/h), pcGECKO suffered a severe proteomic bottleneck, yielding a negligible growth rate (μ ≈ 0.0044 1/h). This result indicates a critical proteomic bottleneck, where the measured enzyme concentrations are insufficient to sustain the high metabolic fluxes required for rapid growth and the onset of the Crabtree effect. Following the standardized protocols suggested in GECKO 3.0 (Chen et al., 2024) to resolve such over-constrained states, two primary correction strategies were evaluated. First, an automated relaxation of turnover numbers, using the sensitivityTuning function, was performed; however, this iteration failed to produce any significant increase in growth. Consequently, a second strategy, proteome mass relaxation via the flexibilizeEnzConcs function, was implemented to force a target empirical growth rate of 0.4 1/h (Van Hoek et al., 1998). While this approach achieved feasibility, it did so at the cost of biological fidelity. Reaching the target growth required the artificial inflation of 74 enzyme mass bounds, often exceeding their original experimental values by several orders of magnitude (Supplementary file 2, Figure S1). This underscores a critical limitation of conventional pcGECKO models: when faced with environmental shifts or metabolic stress, they rely on data manipulation (i.e., overriding empirical evidence) to achieve convergence.

By contrast, the HyTT framework demonstrated superior predictive performance without requiring any artificial relaxation of parameters or data. By utilizing its penalty-driven architecture and an expanded modeled proteome fraction (f = 0.71), HyTT predicted reallocation of resources to resolve the metabolic demand. At the same high-glucose input, HyTT successfully predicted a high growth rate (μ = 0.375 1/h) while accurately capturing the Crabtree effect, with predicted ethanol production and oxygen uptake rates of 10.19 and 5.83 mmol/gDW/h, respectively. Unlike the fixed constraints of pcGECKO, the sequence-derived penalties included in HyTT allowed the model to dynamically partition its capacity, shifting 24.0% of the active proteome mass to the ribosomal machinery and 46.9% to metabolic enzymes under this high-flux condition.

To systematically validate whether this reallocation genuinely reflects biological reality, we benchmarked the absolute molecular abundances predicted by the models against *in vivo* experimental multi-omics datasets at the reference state (Figure 1, glucose uptake = 1.071 mmol/gDW/h, corresponding to the experimental μ = 0.1 1/h condition from Lahtvee et al. (Lahtvee et al., 2017)). Crucially, while both the Static pcGECKO model and the MARS component of HyTT utilized this exact dataset for constraint formulation and initial training, their structural performances differed significantly. The Basic GECKO model and the Static pcGECKO model yielded global proteomic Pearson correlations of r = 0.278 (Spearman ρ = 0.265) and r = 0.283 (Spearman ρ = 0.269), respectively (Figure 1C, 1D). By applying the sequence-driven MARS constraints, the HyTT model significantly improved this predictive fidelity, achieving a remarkable global correlation of r = 0.501 and a Spearman rank correlation of ρ = 0.550 (Figure 1A) (validation of the predictive power of HyTT across unseen environmental conditions is subsequently evaluated in Section 3.3). As a result, HyTT predicts decoupling of translation from metabolism. While standard frameworks treat the cellular proteome as an undifferentiated mass of enzymes, HyTT explicitly distinguishes between metabolic components (r = 0.468) and the translational machinery (r = 0.541).

**Figure 1.**
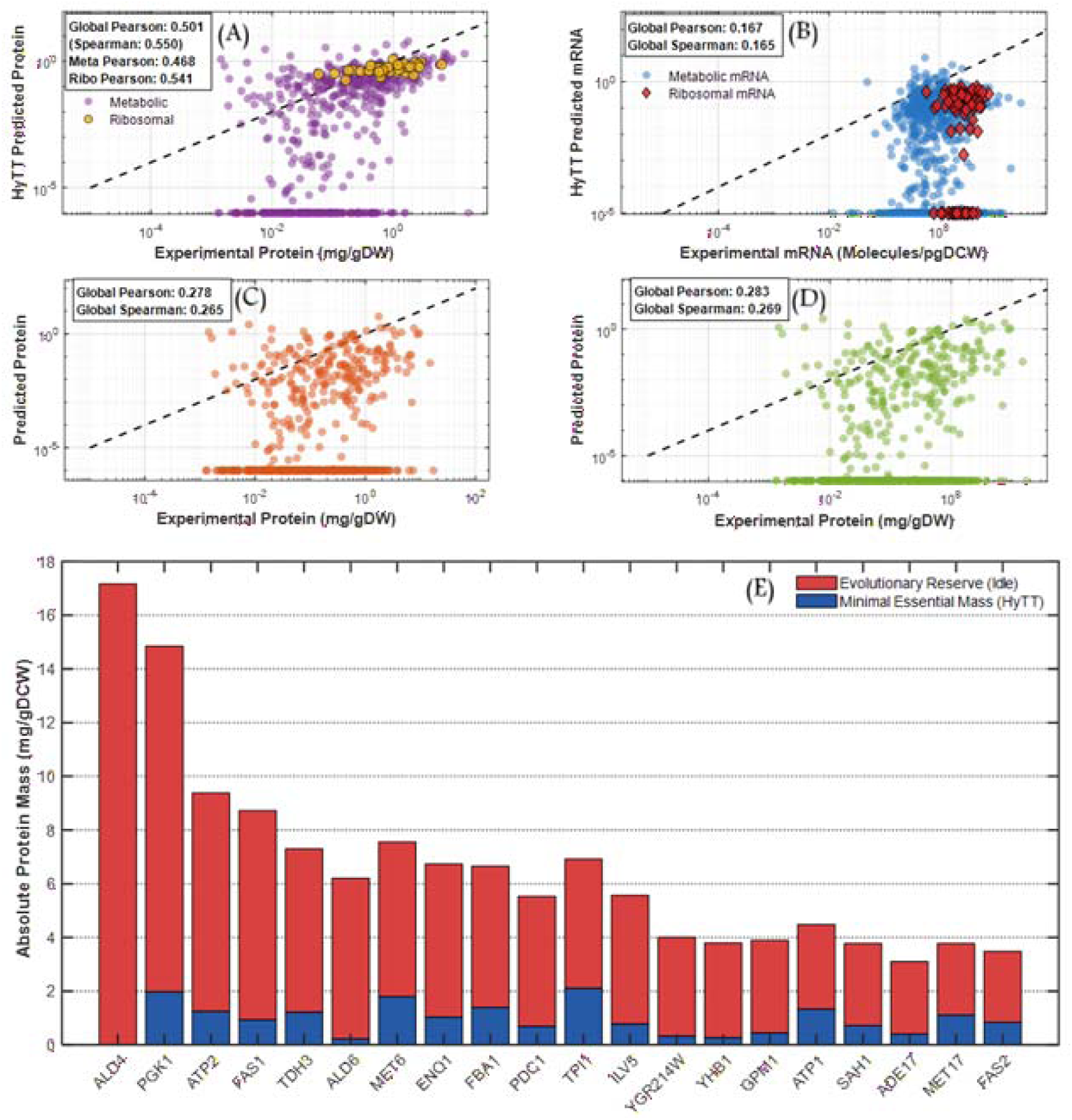
Performance of HyTT with glucose as a carbon source. The panels depict the performance of predicted molecular phenotypes against measured values from omics technologies at a reference state of (Glucose = 1.071 mmol/gDW/h) (Lahtvee et al., 2017). (A) Absolute proteome allocation. (B) Sequence-driven predictions of mRNA. Predicted absolute proteome allocation by (C) Basic GECKO and (D) Static pcGECKO. Pearson and Spearman correlation coefficients were calculated on log10-transformed absolute abundances (g/gDCW). (E) *Ab initio* mapping of the evolutionary reserve proteome in *S. cerevisiae*. The stacked bar chart illustrates the absolute mass distribution (mg/gDCW) of the top 20 proteins with the highest idle capacity. Deep blue sections represent the minimal catalytically essential mass predicted autonomously by the HyTT framework. Red sections indicate the reserve (idle capacity) actively stockpiled by the cell, derived by subtracting the *ab initio* prediction from the total experimental abundance (Lahtvee et al., 2017).

Furthermore, HyTT uniquely extends this predictive capability to the transcriptome (Figure 1B), predicting experimental mRNA abundances with a global Pearson correlation of r = 0.155 and a Spearman correlation of ρ = 0.194. It is important to note that this systemic underprediction of the transcriptome is a well-documented characteristic of parsimony-based optimization, similarly inherent to canonical *E. coli* ME-models (O’Brien et al., 2013). While optimization frameworks apply parsimony across all macromolecules, the resulting predictive gap is substantially wider for the transcriptome than for the proteomic layer (r = 0.501). As detailed below, this disparity stems from a fundamental divergence between strict mathematical optimality and the biological reality of how cells differentially manage transcriptomic versus proteomic reserves, as well as the algorithmic training focus of the MARS model. Notably, the model autonomously captures the massive transcriptional investment required during the growth phase; ribosomal transcripts (Figure 1B, red dots) are correctly clustered in the high-abundance top-right quadrant, effectively mirroring their biological essentiality without any condition-specific transcriptomic input data.

#### Mapping the reserve proteome

While the HyTT framework strictly calculates the minimal catalytically essential proteome, comparing these *ab initio* predictions against empirical absolute abundances (at the reference state of μ = 0.1 1/h from (Lahtvee et al., 2017)) reveals condition-specific idle capacities. By subtracting the predicted essential mass from the experimental reality, we successfully mapped the reserve proteome of *S. cerevisiae*.

The algorithm uncovered a highly strategic, non-random allocation of surplus resources. Whilst the model identified 295 proteins with idle capacity, the top 20 proteins alone account for 46.7% of the total reserve mass (Figure 1E). Aldehyde dehydrogenases (ALD4 and ALD6), functioning as an anticipatory detoxifying reserve against abrupt Crabtree-induced acetaldehyde accumulation, were found as the most prominent buffer. Metabolic flux validation confirmed that at the reference state, *S. cerevisiae* operates in a fully respiratory mode where pyruvate is strictly channeled into the TCA cycle; consequently, the lack of acetaldehyde production explains why this massive ALD pool remains structurally idle *in vivo*. Furthermore, the cell actively stores massive reserves of core glycolytic enzymes (e.g., PGK1, TDH3, ENO1, and PDC1). This pre-translated glycolytic buffer acts as a primed bioenergetic reserve, allowing the yeast to instantly trigger overflow metabolism upon glucose upshifts without enduring the transcriptional and translational delays, and the associated translational costs, of synthesizing new proteome-heavy enzymatic complexes.

#### Understanding the parsimony gap and breaking mathematical symmetry

While the HyTT framework demonstrates high fidelity in capturing global regulatory shifts, examining its absolute mass predictions at the reference state unveils a fundamental characteristic of parsimony-based optimization. At μ = 0.1 1/h, the model allocates only ∼42.7% of the total proteome capacity to satisfy the metabolic demand, representing an expected underprediction relative to the empirical ∼71% saturation (a ∼40% idle reserve). This gap is even more pronounced at the transcriptomic layer; the model calculates a strict minimal mRNA mass of 0.000628 g/gDW (constituting ∼1.03% of total RNA). When compared to the experimental baseline of 0.001202 g/gDW, it becomes evident that the cell actively stockpiles a massive ∼48% of its transcriptome as a completely idle reserve.

This systematic underprediction stems from a fundamental divergence between the mathematical optimality and biological reality. First, from a biophysical perspective, the L1-norm parsimony constraint forces the MILP solver to calculate the strict minimal catalytic requirement to sustain steady-state flux, inherently filtering out the anticipatory buffering pool. Biologically, because mRNA synthesis is energetically inexpensive and occupies negligible physical space compared to massive protein complexes, cells maintain a proportionally larger transcriptomic reserve (∼48%) than a proteomic reserve (∼40%) to remain primed for rapid environmental adaptation. Crucially, as shown above, this parsimony gap does not result in unrealistic predictions that indiscriminately shrink the active network; rather, it represents a highly targeted biological reality, where nearly half of the unpredicted mass (46.7%) is deliberately localized to just 20 preparatory enzymes. Second, from an algorithmic perspective, the MARS model was explicitly trained to predict the translational burden at the proteomic level, not transcriptomic abundance. Therefore, the framework’s ability to predict mRNA allocation is entirely an emergent property enforced by the internal biophysical template lock, naturally explaining the higher predictive fidelity observed for the trained proteomic layer (r = 0.501) compared to the emergent transcriptomic layer (r = 0.155).

By isolating this strict minimal baseline, HyTT preserves highly robust relative topological proportions and effectively resolves a pervasive challenge in constraint-based modeling: the artifact of alternative optima. In highly symmetric or flat objective landscapes, such as those in Basic GECKO and Static pcGECKO, models prematurely collapse into an identical, highly compressed metabolic routing strategy under condition-specific constraints. They severely fail to capture the long tail of true *in vivo* proteomic diversity (Supplementary file2, Figure S2).

The HyTT framework fundamentally breaks this mathematical symmetry *ab initio*. By penalizing the translational burden with explicit sequence features, every reaction carries a highly unique price tag. As demonstrated in Supplementary file 2, Figure S2, under glucose limitation (glucose = 1.071 mmol/gDW/h), this rugged objective landscape allows HyTT to maintain an extensive basal proteomic diversity (expressing >600 functional proteins), avoiding the artificial network collapse seen in standard models. Crucially, under the severe spatial constraints of the Crabtree effect (glucose = 11.1 mmol/gDW/h), HyTT autonomously restructures this hierarchy. To accommodate the massive physical expansion of the ribosomal machinery, visually evident by the clustering of ribosomal components at the absolute highest abundance ranks, the model accurately predicts a targeted pruning of the low-abundance tail, sacrificing non-essential diversity to sustain maximum growth.

Ultimately, this realistic capturing of the hidden translational burden makes the HyTT framework exceptionally well-suited for modeling the production of recombinant or heterologous proteins, which is studied in section 3.4. Furthermore, by recognizing the true metabolic costs of sub-optimal codon usage, specific amino acid demands, and mRNA thermodynamic barriers, HyTT accurately predicts the systemic resource re-allocation imposed by synthetic constructs, providing a powerful tool for rational strain design and metabolic engineering.

### 3.3. Comprehensive chemostat validation and the onset of the Crabtree effect

The HyTT framework provides a robust prediction of the specific growth rate, oxygen consumption, and the production rates of CO_2_ and ethanol when compared against experimental chemostat data (Van Hoek et al., 1998) (Figure 2). The model autonomously identifies the onset of the Crabtree effect at a specific growth rate of µ = 0.35 1/h, precisely where the model predicts a topological breaking point: a sharp decline in the metabolic proteome fraction to accommodate the expanding translational machinery. While this theoretical threshold is slightly higher than the experimentally estimated critical dilution rate of 0.28 to 0.30 1/h (Van Hoek et al., 1998), it successfully captures the fundamental metabolic phase shift derived purely from ab initio biophysical constraints.

**Figure 2.**
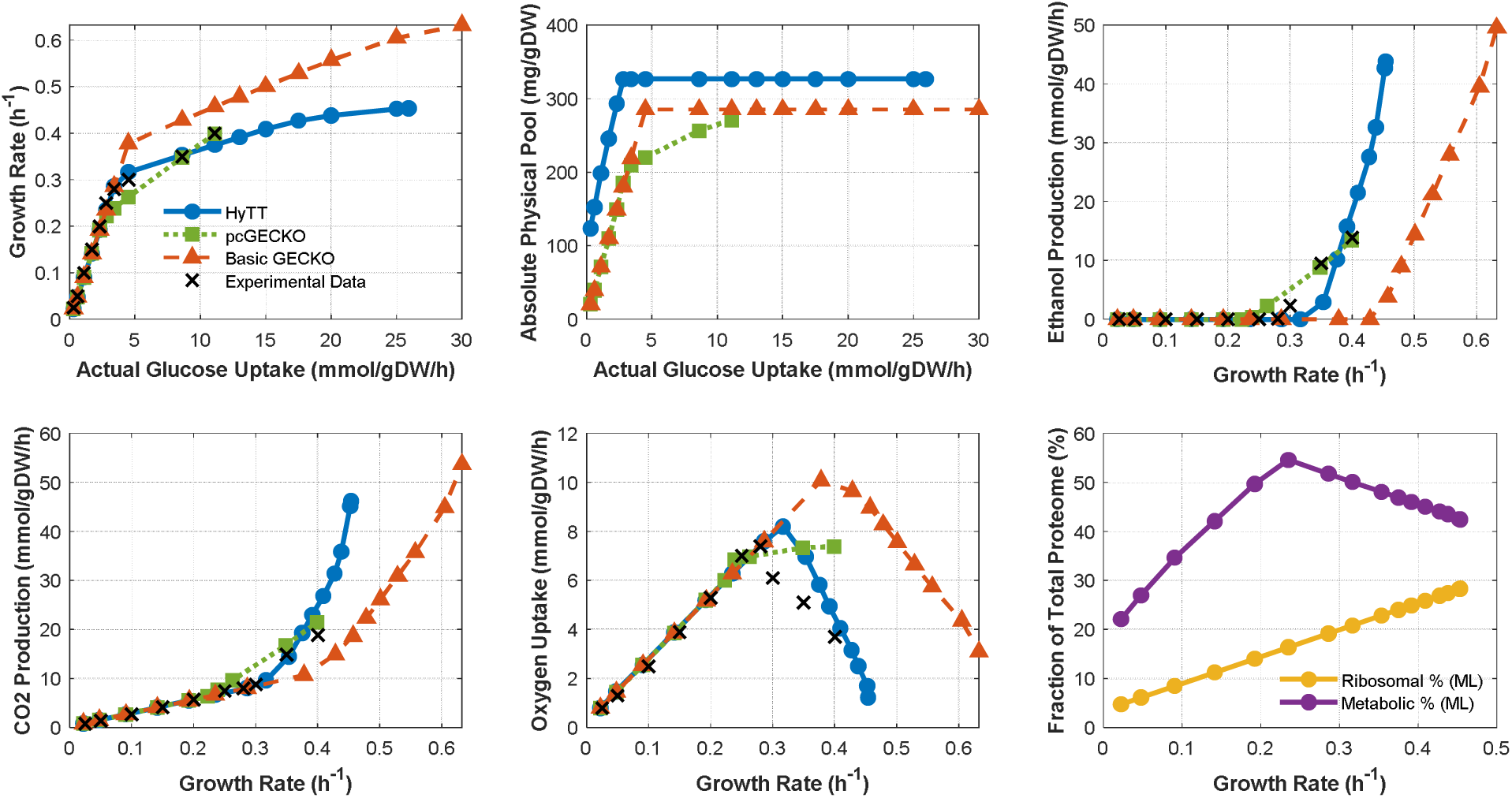
Validation of the HyTT framework against standard models across a glucose-limited chemostat gradient. Simulated macroscopic fluxes, specific growth rate, ethanol production, oxygen consumption, and CO_2_ evolution, and required total proteome by HyTT, Basic GECKO, and pcGECKO are plotted against experimental steady-state data (Van Hoek et al., 1998). The x-axis (Actual Glucose Uptake) reflects the biologically feasible flux bounded autonomously by internal proteomic constraints rather than external theoretical limits.

In contrast, the Basic GECKO model, while performing well under strict glucose-limiting conditions, shows significant divergence upon reaching its initial active proteome capacity limit at a glucose uptake rate of ∼2.8 mmol/gDW/h. Beyond this threshold, Basic GECKO suffers from a notable overprediction of the growth rate and an underprediction of CO_2_ production, delaying its estimated Crabtree onset until µ = 0.46 1/h. The pcGECKO model generally offers a good fit for growth and ethanol profiles, although it estimates an early onset of the Crabtree effect at µ = 0.24 1/h and fails to capture the characteristic decline in oxygen consumption at high growth rates. This failure stems from the model reaching its global proteome ceiling at an artificially high glucose uptake of 11.1 mmol/gDW/h, achieved only after forcing a target maximum growth rate of 0.4 1/h via automated data relaxation (flexibilizeEnzConcs) (Sánchez et al., 2017). This delay in reaching the true proteomic global ceiling in pcGECKO is an artifact of highly restrictive local proteomic constraints on specific enzymes rather than the total physical proteome limit.

Notably, both the HyTT and Basic GECKO models correctly identify that the active catalytic proteome capacity is initially saturated at a much lower glucose uptake rate of ∼2.8 mmol/gDW/h. It is important to highlight the biological sequence of events here: prior to this saturation point, the environmental glucose uptake rate remains the sole limiting factor for cellular growth. However, once this initial catalytic ceiling is hit, the cell enters a strict reallocation regime, where any further increase in growth rate is entirely governed by the internal optimization of proteome mass.

Furthermore, the HyTT framework reveals that as the growth rate increases, the cell’s demand for massive ribosomal complexes, necessary to sustain faster translation rates, can only be met by forcibly displacing the proteome mass allocated to metabolic enzymes (Figure 2). This structural breaking point coincides exactly with the Crabtree onset at µ = 0.35 1/h. This physical trade-off perfectly mirrors recent quantitative mass spectrometry studies. For instance, Metzl-Raz et al. (Metzl-Raz et al., 2017) experimentally demonstrated that at high growth rates, the translation of ribosomal proteins competes with the production of non-ribosomal proteins within a finite cellular volume.

To cope with this severe spatial restriction, the cell is forced to redistribute its metabolic fluxes. As experimentally validated by (Basan et al., 2015), overflow metabolism is a programmed global response used by cells to balance the conflicting proteomic demands of energy biogenesis and biomass synthesis. Because the proteomic cost of generating ATP via respiration is significantly heavier than via fermentation, the cell sacrifices respiratory efficiency to free up physical space for the expanding ribosomal sector. Consequently, the simulated cell in HyTT is incapable of utilizing an unconstrained environmental glucose supply and faces an absolute translational bottleneck, capping the maximum achievable growth rate at µ = 0.45 1/h. This autonomous limitation aligns seamlessly with the experimentally reported maximum aerobic growth rate for *S. cerevisiae* (µ_max_ = 0.40 to 0.45 1/h) (Van Hoek et al., 1998; Verduyn et al., 1990), confirming that sequence-dependent ribosomal expansion and strict physical assembly fundamentally define the ultimate limits of cellular growth and inherently trigger the Crabtree effect.

### 3.4. Systemic proteome burden and sequence-driven reallocation

#### Intensification of the Crabtree effect under GFP-induced protein burden

Building upon the strictly proteome-limited physiological state established at a high baseline glucose uptake rate (11.1 mmol/gDW/h), we sought to evaluate the predictive capabilities and robustness of HyTT under conditions of severe metabolic engineering stress. To simulate an acute and systemic proteome burden, the heterologous production of 15% GFP was integrated into the core metabolic network. Because the wild-type *in silico* cell at this specific carbon influx rate is already operating at its global physical ceiling (achieving a 71% active proteome saturation), the sudden 15% reduction in available endogenous allocation space acts as a severe restriction, triggering a profound, system-wide reallocation response.

Under this extreme constraint, HyTT autonomously predicts a macroscopic metabolic phase shift (Figure 3A). To survive within the artificially restricted proteomic volume, the cell is forced to dismantle its bulky, high-molecular-weight respiratory machinery. Specifically, the model predicts that the specific oxygen uptake rate drops from 5.83 to 3.95 mmol/gDW/h. To fully compensate for the consequent loss in highly efficient respiratory ATP generation, the cell dynamically hyper-activates its highly proteome-efficient glycolytic and fermentative pathways, causing the specific ethanol production rate to surge from 10.19 to 13.98 mmol/gDW/h. This emergent behavior serves as an *ab initio* evidence for overflow metabolism as fermentative pathways produce less ATP per carbon unit but require significantly less physical proteome mass per ATP generated; the cell sacrifices high-yield respiratory carbon conversion. It strategically trades carbon efficiency for proteome efficiency, maximizing absolute bioenergetic output within a severely restricted proteomic capacity (Basan et al., 2015).

**Figure 3.**
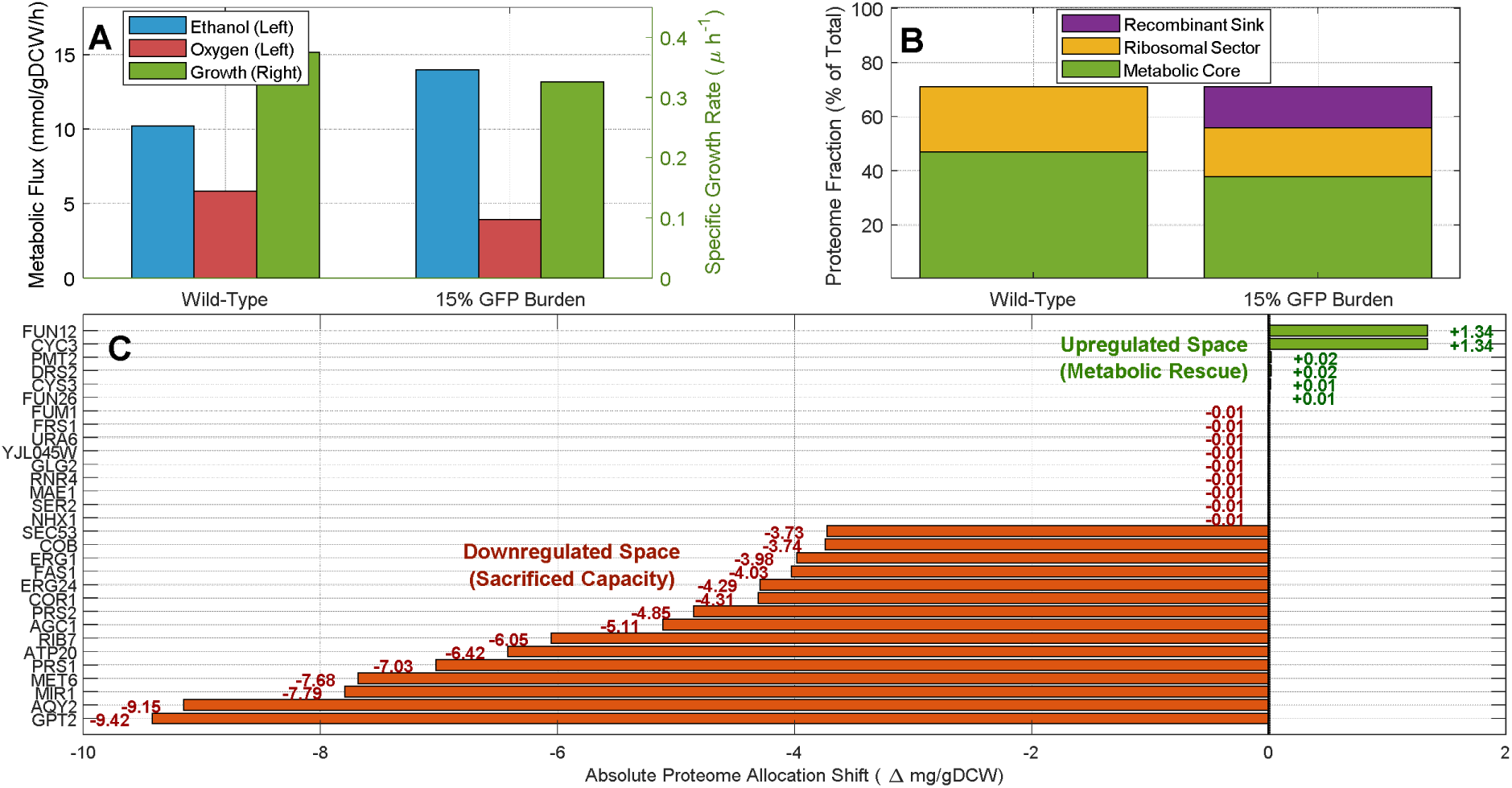
Systemic metabolic and proteomic reallocation under GFP-induced burden predicted by HyTT. **(A)** Macroscopic flux and growth rate retardation under a 15% heterologous GFP burden. **(B)** Dynamic proteome reallocation. **(C)** Metabolic re-routing. Horizontal bars represent the absolute shift in proteome allocation (Δ mg/gDW) for down-regulated (red bars) and up-regulated (green bars) enzymes.

#### Ribosomal reduction as a response strategy to metabolic burden

Crucially, HyTT uncovers the mechanistic logic driving this systemic adaptation by capturing the fundamental biological trade-off between energy demand and translational capacity. As the 15% GFP burden collides with the hard spatial ceiling, the MILP solver mathematically dictates a necessary proportional reduction in the specific growth rate (a ∼13% reduction, dropping from 0.375 to 0.326 1/h).

This ∼13% reduction in the specific growth rate in response to a 15% heterologous burden is not a random numerical penalty; rather, it autonomously recapitulates the well-established linear growth law (Scott et al., 2010). As the functional proteome capacity is restricted by 15%, the maximum achievable growth rate drops proportionally (Metzl-Raz et al., 2017). The fact that the growth retardation (∼13%) is slightly less severe than the exact volumetric burden (15%) highlights the model’s ability to capture biological resilience, specifically the cell’s utilization of its pre-existing idle reserve proteome to partially buffer the metabolic burden, a proportional behavior exactly consistent with *in vivo* observations of yeast overexpressing fluorescent proteins (Eguchi et al., 2018; Kafri et al., 2016).

By achieving this slower growth rate, the cell’s inherent biophysical demand for actively translating ribosomes decreases proportionally. Specifically, HyTT predicts a dramatic and highly targeted contraction of the ribosomal machinery, shrinking its absolute mass fraction from 24.0% of the total proteome in the wild-type state down to 18.1% under the recombinant stress (Figure 3B). This targeted liberation of proteomic real estate serves a dual evolutionary purpose. First, it acts as a primary physiological response to mitigate the heterologous burden. Second, it explicitly frees up the critical intracellular space required to accommodate the expanded glycolytic enzymes needed to sustain emergency fermentative ATP generation. By dynamically coupling these sequence-specific translational costs to absolute spatial constraints, HyTT mathematically confirms that growth retardation is an active survival strategy employed to balance the expanding metabolic core against the heavily sequestering ribosomal sector (Metzl-Raz et al., 2017).

#### Non-linear metabolic redistribution and proteome economy

Whilst macroscopic flux analysis clearly demonstrates a shift towards fermentative metabolism, a higher-resolution interrogation of the endogenous proteome (Figure 3C) reveals that the 15% spatial burden does not compress the metabolic network uniformly. Instead, the HyTT framework predicts a highly targeted, non-linear metabolic redistribution. While the applied L1-norm minimization ensures basic network parsimony, identifying the minimal essential flux adjustments required to support the new stress-induced growth rate, the fundamental driver of this metabolic shift is the constrained proteomic capacity and HyTT’s sequence-derived translational penalties, which actively force the prioritization of the proteome-compact pathways. This redistribution is characterized by the dismantling of massive multimeric complexes and the strategic upregulation of proteome-efficient, high-turnover enzymes to optimize global network bioenergetics. A striking example of this proteomic optimization occurs in the systematic down-regulation of the space-consuming multimeric complexes required for oxidative phosphorylation and heavy biosynthesis. Core subunits of the Cytochrome bc1 complex (e.g., COR1, COB) and the massive F1F0-ATP synthase (e.g., ATP20), alongside spatially expensive sinks like Fatty Acid Synthase (FAS1) and key nucleotide synthesis complexes (e.g., PRS1, PRS2), are heavily penalized and sacrificed by the solver (Figure 3C, red bars).

Simultaneously, to maintain essential metabolic fluxes and bioenergetic homeostasis within the restricted proteomic capacity, the model upregulates highly compact, high-turnover alternatives (Figure 3C, green bars). A prominent example is the upregulation of Triosephosphate isomerase (TPI1) and Pyruvate decarboxylase (PDC1). Because enzymes like TPI1 operate near the catalytic diffusion limit (exhibiting phenomenally high kcat values), they allow the cell to maximize carbon throughput and ATP yield while occupying only a minuscule fraction of the restricted protein mass. By autonomously prioritizing enzymes with extreme turnover numbers and lower sequence-dependent translational costs, the HyTT model provides rigorous quantitative proof that metabolic re-routing under a heterologous protein burden is fundamentally driven by strict principles of spatial economy.

#### Ribosomal paralog switching sequence-driven translational optimization

Beyond purely metabolic realignments, the integration of strict 80S stoichiometric constraints alongside the sequence-aware MARS equations unveiled a deeper layer of systemic adaptation. Unlike the flexible metabolic network, where the cell can actively downregulate physically massive enzyme complexes to conserve space, the cellular translation machinery is governed by rigid 1:1 and 2:1 stoichiometric assembly rules. The cell cannot simply omit large ribosomal subunits to save mass without completely halting global translation. Furthermore, alternative ribosomal paralogs (e.g., RPS6A and RPS6B) possess functionally identical physical molecular weights, thereby offering absolutely zero macroscopic mass savings upon switching.

Faced with this rigid assembly constraint, the HyTT framework reveals a dual-layered evolutionary optimization strategy. While the cell adapts to the proteome burden in the metabolic sector by minimizing mass allocation (downregulating proteome-expensive enzyme complexes), it mitigates stress within the ribosomal sector by explicitly minimizing sequence-dependent translational costs. Under the severe translational burden induced by the heavy demand of recombinant GFP transcripts, the model systematically represses kinetically expensive paralogs characterized by higher translational costs in the MARS formulation (e.g., RPS6A, RPS8A, and RPL2A), as shown in Figure 4. Concurrently, the model upregulates translationally optimized, cheaper paralogs possessing superior sequence architectures (e.g., RPS6B [CAI: 0.903 vs 0.892], RPS8B [CAI: 0.912 vs 0.900], and RPL2B [CAI: 0.909 vs 0.902]).

**Figure 4.**
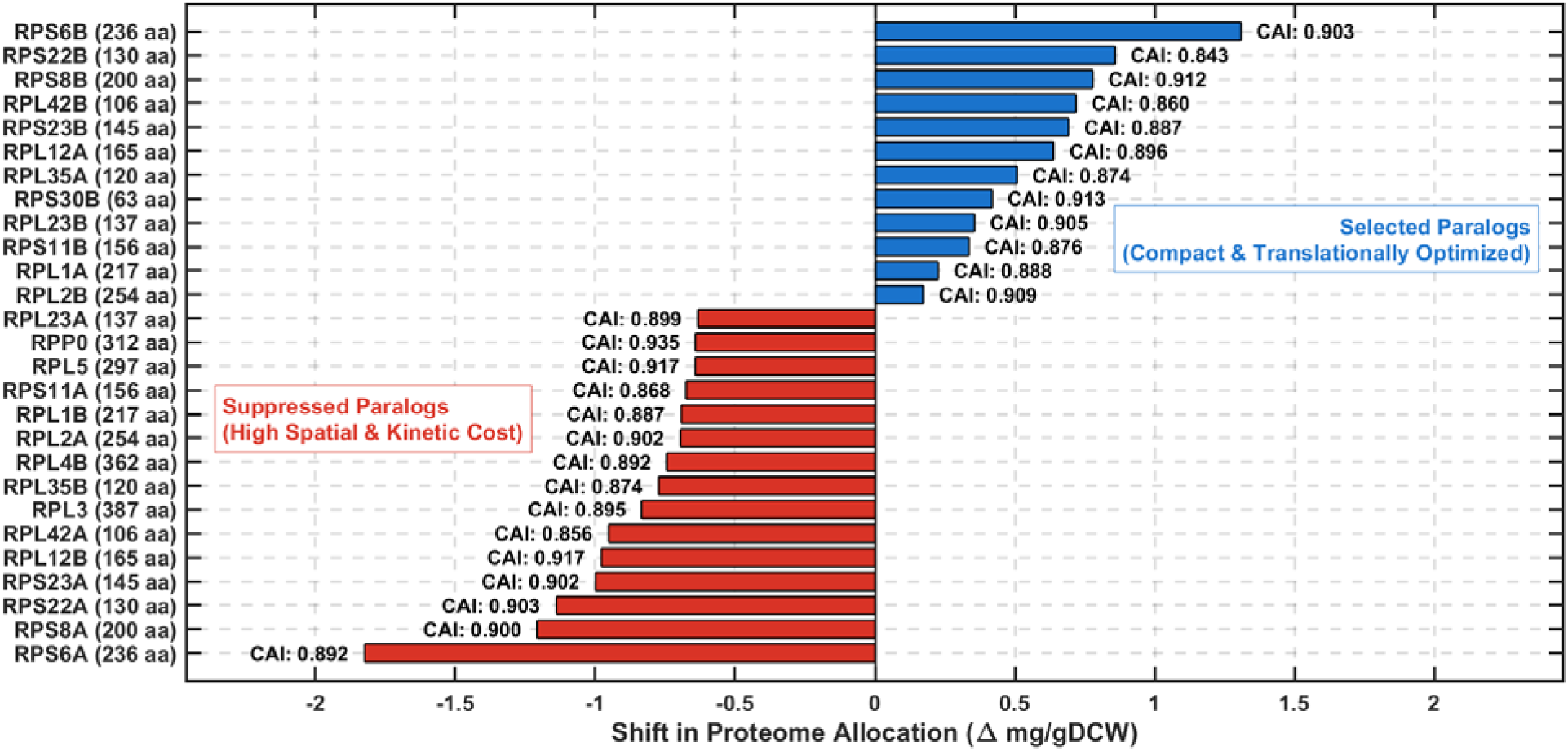
Targeted ribosomal paralog switching under a 15% recombinant burden. Horizontal bars represent the absolute shift in proteome allocation (Δ mg/gDCW) for specific ribosomal proteins. Rather than a uniform downregulation of the ribosomal sector, the model downregulates translationally expensive paralogs (red bars) in favor of their more efficient counterparts (blue bars). This sequence-driven reallocation minimizes the translational cost of assembling the 3 MDa 80S complex under recombinant stress.

Because these paired paralogs fulfill the exact same structural subunit role within the 80S complex but carry distinct synthesis costs, this mathematically autonomous switching is profoundly significant. It successfully mirrors the biological reality of ribosomal heterogeneity. Recent advanced translatome and proteome profiling studies have empirically demonstrated that wild-type yeast actively alters the stoichiometric ratios of its ribosomal paralogs in response to severe environmental stress, shifting from major to minor paralog expression to remodel ribosome composition and preserve translational fidelity (Ghulam et al., 2020). By successfully predicting this targeted paralog switching purely from ab initio sequence constraints, HyTT demonstrates that the cell mitigates the burden of 80S assembly by explicitly minimizing sequence-dependent elongation costs when global proteomic space is critically constrained.

## 4. Conclusion

The Hybrid Translation-Transcription (HyTT) framework introduces a mechanistic-statistical paradigm to resolve the intractable parameterization bottlenecks of traditional constraint-based models. By integrating MARS with the rigorous stoichiometric backbone of enzyme-constrained networks, HyTT successfully captures the highly complex, non-linear hidden patterns governing sequence-specific translation efficiencies. Instead of explicitly simulating every microscopic biochemical transaction via non-linear differential equations, HyTT seamlessly injects these translational complexities directly into the optimization landscape as empirical, ab initio MILP constraints. This mathematical duality enables HyTT to preserve the structural realism of macromolecular expressions while achieving unprecedented computational agility.

A core principle underlying the HyTT framework is the dynamic interplay between global macroscopic boundaries and sequence-specific microscopic constraints. The total intracellular proteome pool and carbon uptake flux act as overarching macro-constraints, defining the absolute physical and bioenergetic ceiling of the cell. Crucially, rather than relying on condition-specific *omics* data or rigid empirical upper bounds, HyTT enforces a single, unified macro-capacity pool (f_pool_ = 0.71). Within this global space, the MARS-derived equations act as micro-constraints, assigning a precise translational cost to each gene based on its unique sequence architecture. By employing L1-norm parsimony across this unified pool, the MILP solver autonomously partitions resources between the active metabolic core and the 80S ribosomal machinery. This completely eliminates a foundational vulnerability of conventional models: the reliance on fixed metabolic proteome fractions (*f*-factors) that artificially paralyze the network under stress. Unlike these rigid frameworks, HyTT enables a dynamic, condition-responsive trade-off between metabolic flux and translational demand. By allowing the optimization algorithm to resolve the tension between strict macro-capacity limits and sequence-dependent micro-costs, HyTT directly links genomic architecture to systems-level resource allocation, providing an unbiased, *ab initio* window into true cellular economy.

HyTT proves particularly powerful for decoding the systemic stress of recombinant protein production. By locking this exogenous burden to the biomass objective, the model forces a proportional contraction of the endogenous proteomic space, mechanistically capturing the fundamental trade-off between host growth and recombinant yield. Cellular adaptation to this stress manifests not as a uniform reduction, but as a highly targeted metabolic redistribution. By mapping the endogenous genome’s sequence characteristics to their systemic translational costs, HyTT exposes the true metabolic burden of recombinant expression. For metabolic engineers, this highlights that optimizing the heterologous construct alone is fundamentally insufficient if the host’s entire endogenous network remains constrained by inherent sequence-dependent limitations and global capacity bounds. Ultimately, HyTT provides a sequence-driven, computationally tractable platform for decoding dynamic proteome reallocation without requiring condition-specific omics inputs from the investigated scenario, offering a robust tool for rational strain design and next-generation metabolic engineering.

## Acknowledgements

This work was supported by a research fellowship from the Alexander von Humboldt Foundation awarded to E.M.

## Data and code availability

The HyTT scripts developed for this study are openly available in the GitHub repository at https://github.com/ehsanmotamedian/HyTT. The comprehensive datasets supporting the conclusions of this article, including the extracted biophysical features and the MARS-derived parameters for the mapped network genes, are provided in the Supplementary Material. The base genome-scale metabolic model (ecYeastGEM) and the experimental multi-omics datasets utilized for model training and validation are derived from previously published studies and are appropriately cited within the text.

