## Supplementary file 2 for "A hybrid machine learning and enzyme-constrained metabolic model for *ab initio* prediction of proteome reallocation"

**A hybrid machine learning and enzyme-constrained metabolic model for *ab initio* prediction of proteome reallocation**

Ehsan Motamedian and Zoran Nikoloski

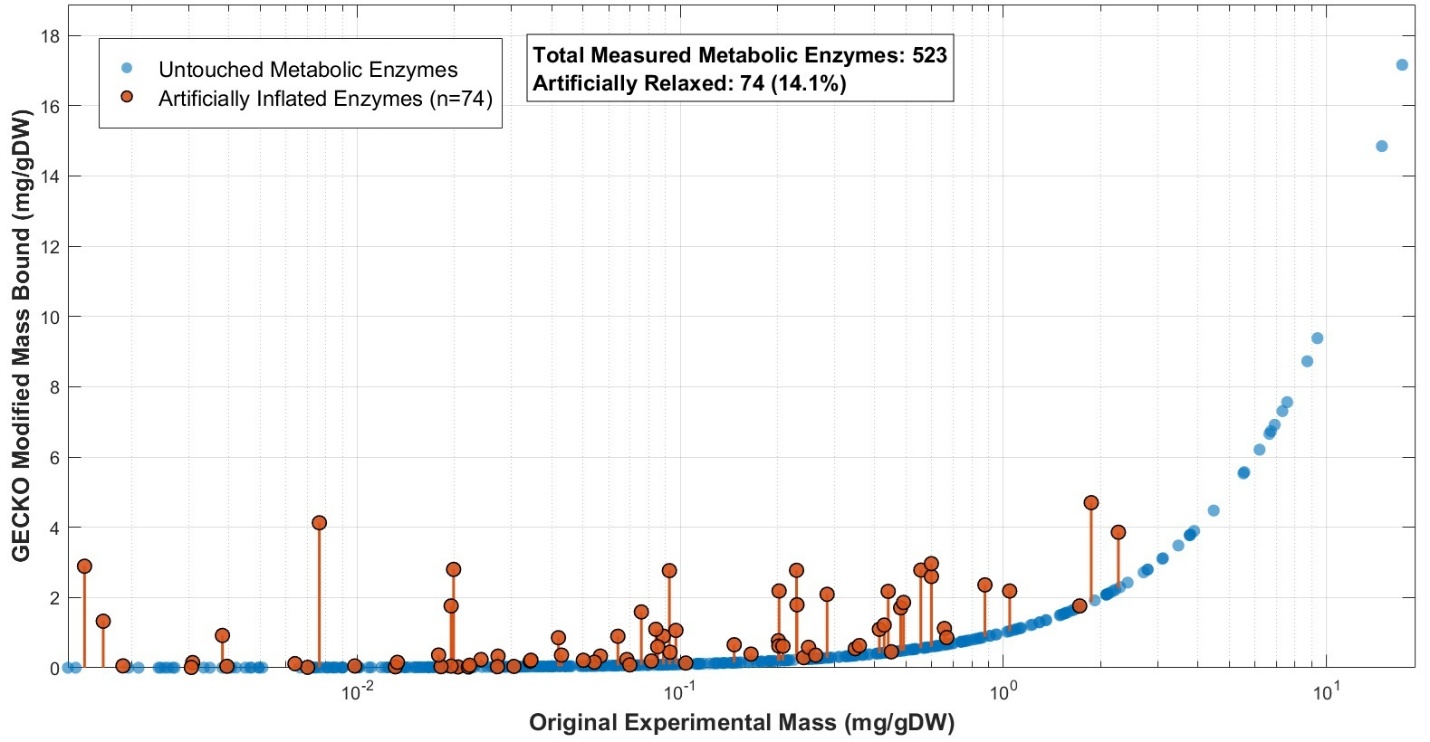

**Figure S1.** Proteomic data manipulation required by the classic pcGECKO model to achieve feasibility at high glucose uptake rates. The scatter plot highlights the subset of enzymes whose experimental mass bounds were artificially inflated (flexibilized) to force mathematical convergence.

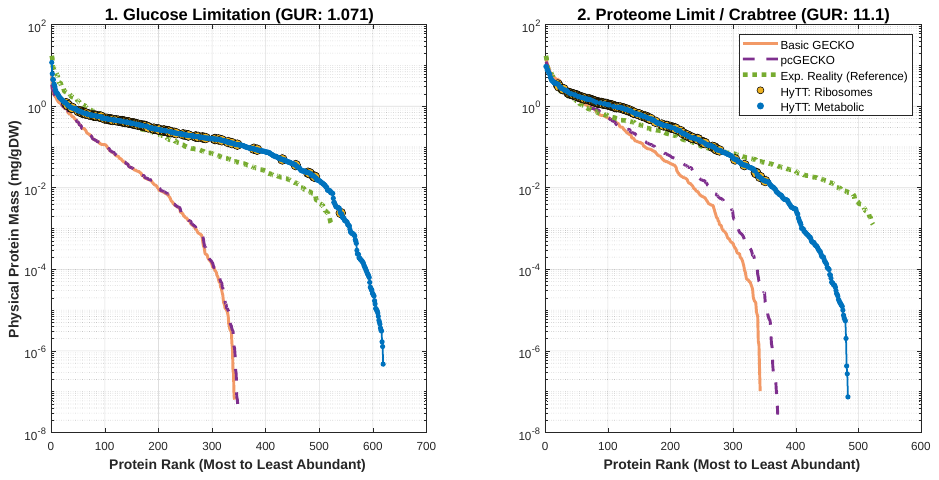

**Supplementary Figure S2. Dynamic proteome allocation and rank-abundance profiles across distinct physiological states.** The panels compare the predicted physical protein mass distributions of the Basic GECKO (orange), Static pcGECKO (purple dashed), and HyTT (blue for metabolic, gold for ribosomal) models against *in vivo* experimental reality (green dotted line). All models were subjected to an identical strict parsimony constraint (minimization of total protein pool usage) to evaluate fundamental objective landscape symmetry. **(1)** Under strict glucose limitation (glucose uptake = 1.071 mmol/gDW/h), standard models perfectly overlap and artificially collapse the network into a highly compressed routing strategy. In contrast, HyTT mathematically breaks this symmetry via sequence-driven translational penalties, successfully capturing the long tail of basal proteomic diversity (>600 expressed proteins). **(2)** At the global proteome limit (glucose uptake = 11.1 mmol/gDW/h), the severe spatial constraint forces a global topological shift. HyTT autonomously predicts the pruning of low-abundance non-essential proteins to physically accommodate the massive expansion of the translational machinery, clearly visualized by the aggressive upward and leftward shift of ribosomal components (gold circles) into the highest abundance ranks.

**Supplementary Note S1: Handling Model Feasibility and Sink Reactions**

Furthermore, to maintain strict steady-state conditions and prevent mathematical dead-ends during resource reallocation, dedicated sink reactions (sink_mRNA_uniprot and sink_prot_uniprot) were appended for every transcript and protein species. Crucially, these sinks serve distinct computational and biological purposes within the hybrid MILP coupling architecture. While the mRNA sink enables the quantitative expression of transcripts to satisfy the MARS equations, the protein sink functions as an algorithmic escape valve. Because the piecewise-linear MARS equations can mathematically enforce the network to over-synthesize certain enzymes beyond the immediate stoichiometric demands of metabolic fluxes, the unconstrained protein sink prevents computational infeasibility by purging this surplus mass, thereby ensuring robust model convergence under strict coupling constraints. This hierarchical penalty system successfully simulates the continuous energetic maintenance cost of transcript turnover and protein degradation, guiding the solver toward a parsimonious, biologically realistic distribution without artificially choking essential transcription. **Supplementary Note S2: Mathematical and Stoichiometric Implementation of Recombinant GFP Burden**

To rigorously simulate the metabolic stress induced by recombinant protein expression, the synthesis of Green Fluorescent Protein (GFP) was integrated into the HyTT framework not merely as an arbitrary ATP sink, but as a sequence-specific, mass-occupying entity physically coupled to cellular proliferation.

**1. Sequence-Specific Stoichiometry and Energetic Costs**

The *in silico* synthesis reaction (r_GFP_synthesis) was explicitly defined based on the precise amino acid composition of the Enhanced Green Fluorescent Protein (EGFP) variant modeled in this study (length = 239 amino acids, MW ≈ 26.89 kDa). The stoichiometric formulation exactly maps the specific molar requirements of all 20 proteinogenic amino acids withdrawn from the intracellular precursor pools (detailed in Supplementary Table S1).

Furthermore, the thermodynamic cost of translation was rigorously integrated. The model explicitly accounts for the high-energy phosphate bonds—specifically 952 ATP and 476 GTP molecules—utilized during the tRNA aminoacylation phase and by the ribosomal elongation and translocation factors. The summarized stoichiometric equation appended to the standard *S. cerevisiae* stoichiometric matrix (S) is represented as:

∑(n_i_ × Amino Acid_i_) + 952 ATP + 476 GTP → GFP + 237 H_2_O + 952 ADP + 476 GDP + 714 P_i_

*(Note: In the model's standardized nomenclature, energy metabolites were mapped to their respective specific compartmentalized IDs, preserving the explicit biological distinction between ATP and GTP usage during translation).*

Supplementary Table S1: Exact Amino Acid Composition and Stoichiometry for Recombinant GFP Synthesis

| **Network ID** | **Amino Acid** | **3-Letter Code** | **Count (Molecules)** | **Relative Abundance (%)** |
| --- | --- | --- | --- | --- |
| s_1003 | Glycine | Gly | 22 | 9.21% |
| s_1021 | Leucine | Leu | 21 | 8.79% |
| s_1025 | Lysine | Lys | 20 | 8.37% |
| s_1056 | Valine | Val | 18 | 7.53% |
| s_0973 | Aspartate | Asp | 18 | 7.53% |
| s_0991 | Glutamate | Glu | 16 | 6.69% |
| s_1045 | Threonine | Thr | 16 | 6.69% |
| s_0969 | Asparagine | Asn | 13 | 5.44% |
| s_1032 | Phenylalanine | Phe | 12 | 5.02% |
| s_1016 | Isoleucine | Ile | 12 | 5.02% |
| s_1051 | Tyrosine | Tyr | 11 | 4.60% |
| s_1039 | Serine | Ser | 10 | 4.18% |
| s_1035 | Proline | Pro | 10 | 4.18% |
| s_1006 | Histidine | His | 9 | 3.77% |
| s_0955 | Alanine | Ala | 8 | 3.35% |
| s_0999 | Glutamine | Gln | 8 | 3.35% |
| s_1029 | Methionine | Met | 6 | 2.51% |
| s_0965 | Arginine | Arg | 6 | 2.51% |
| s_0981 | Cysteine | Cys | 2 | 0.84% |
| s_1048 | Tryptophan | Trp | 1 | 0.42% |
| **Total** | | | **239** | **100.00%** |

**2. Mathematical Coupling to the Global Proteome Constraint**

To simulate a precise 15% mass burden, the production of GFP had to be dynamically coupled to the specific growth rate (μ). Simply forcing a static flux through the GFP synthesis reaction would violate the steady-state assumption at varying growth rates.

Instead, a pseudo-metabolite (p_GFP_demand) was introduced into the S matrix to act as a mass bridge. The continuous synthesis of GFP was mathematically locked to the core biomass objective function (r_4041), ensuring that for every gram of dry cell weight (gDCW) produced, exactly 15% of the total available proteome capacity (P_total_) is physically sequestered as GFP. This was achieved by modifying the coefficient of the GFP demand metabolite in the biomass reaction:

S_(GFP_demand, Biomass)_ = -(P_total_ × 0.15) / MW_GFP_

**3. Activating the Proteome Squeeze**

Once the recombinant mass was successfully coupled to cellular growth, the physical space available for the endogenous metabolic network had to be proportionally contracted. In the HyTT Mixed-Integer Linear Programming (MILP) architecture, the global proteome bound for endogenous enzymes and ribosomes was dynamically updated to:

∑(v_j_ / (k_cat,j_ × σ)) + [P]_ribo_(μ) ≤ f × P_total_ × (1 - 0.15)

By shrinking the right side of the constraint and forcing the biomass reaction to carry the heavy GFP mass, the MILP solver is forced to autonomously reallocate the remaining 85% of the proteomic space. This formulation ensures that the translational machinery must still actively translate the exogenous GFP, incurring the associated sequence-driven MARS penalties, thereby providing a highly realistic *ab initio* simulation of the *in vivo* protein burden phenomenon.
